## Supplementary material for "An expanding manifold in transmodal regions characterizes adolescent reconfiguration of structural connectome organization": Data S1

| Gene symbol | Name | t | FDR |
|---|---|---|---|
| **CHST9** | carbohydrate (N-acetylgalactosamine 4-0) sulfotransferase 9 | -24.76 | 0.0058 |
| **SHD** | Src homology 2 domain containing transforming protein D | 24.20 | 0.0058 |
| **SPTSSB** | serine palmitoyltransferase, small subunit B | 20.90 | 0.0088 |
| **CPNE9** | copine family member IX | 20.12 | 0.0095 |
| **COL5A1** | collagen, type V, alpha 1 | 18.34 | 0.0097 |
| **POLR2L** | polymerase (RNA) II (DNA directed) polypeptide L, 7.6kDa | -18.46 | 0.0097 |
| **RSPO4** | R-spondin 4 | -18.57 | 0.0097 |
| **COX7A1** | cytochrome c oxidase subunit VIIa polypeptide 1 (muscle) | 17.57 | 0.0100 |
| **FAM20A** | family with sequence similarity 20, member A | 17.77 | 0.0100 |
| **ITPR1** | inositol 1,4,5-trisphosphate receptor, type 1 | 17.39 | 0.0100 |
| **HAPLN4** | hyaluronan and proteoglycan link protein 4 | 16.59 | 0.0116 |
| **GLUD1** | glutamate dehydrogenase 1 | -16.27 | 0.0118 |
| **KCNN3** | potassium intermediate/small conductance calcium-activated channel, subfamily N, | -15.51 | 0.0118 |
| **PPL** | periplakin | 15.24 | 0.0118 |
| **RELL2** | RELT-like 2 | 16.34 | 0.0118 |
| **RGS7** | regulator of G-protein signaling 7 | 15.24 | 0.0118 |
| **SMARCD3** | SWI/SNF related, matrix associated, actin dependent regulator of chromatin, subfamil | -15.52 | 0.0118 |
| **PRSS16** | protease, serine, 16 (thymus) | 14.22 | 0.0132 |
| **SCN1A** | sodium channel, voltage-gated, type I, alpha subunit | 14.23 | 0.0132 |
| **STX19** | syntaxin 19 | 14.50 | 0.0132 |
| **NGB** | neuroglobin | 13.77 | 0.0137 |
| **DDAH2** | dimethylarginine dimethylaminohydrolase 2 | -13.61 | 0.0141 |
| **GPD2** | glycerol-3-phosphate dehydrogenase 2 (mitochondrial) | -13.57 | 0.0141 |
| **EPN3** | epsin 3 | 13.29 | 0.0146 |
| **LRRC38** | leucine rich repeat containing 38 | 13.22 | 0.0146 |
| **FBLN7** | fibulin 7 | 13.06 | 0.0150 |
| **MACROD2** | MACRO domain containing 2 | -13.01 | 0.0151 |
| **NKAIN4** | Na+/K+ transporting ATPase interacting 4 | -12.70 | 0.0160 |
| **CPLX1** | complexin 1 | 12.48 | 0.0164 |
| **EXTL2** | exostoses (multiple)-like 2 | 12.53 | 0.0164 |
| **FLRT3** | fibronectin leucine rich transmembrane protein 3 | 12.48 | 0.0164 |
| **LYPD1** | LY6/PLAUR domain containing 1 | -12.38 | 0.0164 |
| **GLCCI1** | glucocorticoid induced transcript 1 | 12.24 | 0.0165 |
| **LAG3** | lymphocyte-activation gene 3 | 12.06 | 0.0168 |
| **DNAH14** | dynein, axonemal, heavy chain 14 | -11.32 | 0.0170 |
| **KCNAB3** | potassium voltage-gated channel, shaker-related subfamily, beta member 3 | 11.95 | 0.0170 |
| **PREP** | prolyl endopeptidase | 11.82 | 0.0170 |
| **SCN1B** | sodium channel, voltage-gated, type I, beta subunit | 11.67 | 0.0170 |
| **SCRT1** | scratch homolog 1, zinc finger protein (Drosophila) | 11.53 | 0.0170 |
| **SHROOM3** | shroom family member 3 | 11.40 | 0.0170 |
| **SPAG4** | sperm associated antigen 4 | 11.30 | 0.0170 |
| **TDRD1** | tudor domain containing 1 | 11.74 | 0.0170 |
| **TNNC2** | troponin C type 2 (fast) | 11.88 | 0.0170 |
| **TPK1** | thiamin pyrophosphokinase 1 | 11.39 | 0.0170 |
| **FGF18** | fibroblast growth factor 18 | 11.18 | 0.0172 |
| **KLHL13** | kelch-like 13 (Drosophila) | -11.16 | 0.0172 |
| **ASB13** | ankyrin repeat and SOCS box containing 13 | 10.81 | 0.0173 |
| **LINC00473** | long intergenic non-protein coding RNA 473 | 10.85 | 0.0173 |
| **MYO15A** | myosin XVA | 11.01 | 0.0173 |
| **S100A10** | S100 calcium binding protein A10 | -10.92 | 0.0173 |
| **SIX4** | SIX homeobox 4 | 10.79 | 0.0173 |

| | | | |
|---|---|---|---|
| **STAMBPL1** | STAM binding protein-like 1 | 10.87 | 0.0173 |
| **STXBP5L** | syntaxin binding protein 5-like | 10.97 | 0.0173 |
| **SLIT3** | slit homolog 3 (Drosophila) | -10.73 | 0.0177 |
| **GABRA1** | gamma-aminobutyric acid (GABA) A receptor, alpha 1 | 10.66 | 0.0179 |
| **FES** | feline sarcoma oncogene | 10.65 | 0.0179 |
| **MIR31HG** | MIR31 host gene (non-protein coding) | 10.57 | 0.0181 |
| **OSBPL6** | oxysterol binding protein-like 6 | 10.59 | 0.0181 |
| **GABRD** | gamma-aminobutyric acid (GABA) A receptor, delta | 10.53 | 0.0182 |
| **SCAPER** | S-phase cyclin A-associated protein in the ER | -10.46 | 0.0185 |
| **GNG4** | guanine nucleotide binding protein (G protein), gamma 4 | -10.38 | 0.0185 |
| **IL17RD** | interleukin 17 receptor D | -10.37 | 0.0185 |
| **SRPK1** | SRSF protein kinase 1 | 10.38 | 0.0185 |
| **EIF4E1B** | eukaryotic translation initiation factor 4E family member 1B | 10.30 | 0.0187 |
| **MAP3K13** | mitogen-activated protein kinase kinase kinase 13 | 10.30 | 0.0187 |
| **NTSR2** | neurotensin receptor 2 | -10.28 | 0.0187 |
| **CFD** | complement factor D (adipsin) | -10.24 | 0.0188 |
| **UCHL3** | ubiquitin carboxyl-terminal esterase L3 (ubiquitin thiolesterase) | -10.20 | 0.0189 |
| **KCNA2** | potassium voltage-gated channel, shaker-related subfamily, member 2 | 10.13 | 0.0191 |
| **PVALB** | parvalbumin | 10.01 | 0.0195 |
| **UCHL5** | ubiquitin carboxyl-terminal hydrolase L5 | 10.02 | 0.0195 |
| **HIVEP2** | human immunodeficiency virus type I enhancer binding protein 2 | 9.90 | 0.0200 |
| **CADPS2** | Ca++-dependent secretion activator 2 | 9.80 | 0.0202 |
| **CITED2** | Cbp/p300-interacting transactivator, with Glu/Asp-rich carboxy-terminal domain, 2 | 9.78 | 0.0202 |
| **GLS2** | glutaminase 2 (liver, mitochondrial) | 9.82 | 0.0202 |
| **STRBP** | spermatid perinuclear RNA binding protein | 9.81 | 0.0202 |
| **PTPRA** | protein tyrosine phosphatase, receptor type, A | -9.75 | 0.0202 |
| **CCNI** | cyclin I | 9.39 | 0.0210 |
| **DERL1** | derlin 1 | -9.38 | 0.0210 |
| **EIF5A2** | eukaryotic translation initiation factor 5A2 | 9.55 | 0.0210 |
| **KBTBD6** | kelch repeat and BTB (POZ) domain containing 6 | -9.46 | 0.0210 |
| **PDLIM5** | PDZ and LIM domain 5 | -9.41 | 0.0210 |
| **SLC39A14** | solute carrier family 39 (zinc transporter), member 14 | 9.45 | 0.0210 |
| **SYCP2** | synaptonemal complex protein 2 | 9.49 | 0.0210 |
| **TPTE2P6** | transmembrane phosphoinositide 3-phosphatase and tensin homolog 2 pseudogen | 9.57 | 0.0210 |
| **ZNF385B** | zinc finger protein 385B | 9.49 | 0.0210 |
| **MAFB** | v-maf musculoaponeurotic fibrosarcoma oncogene homolog B (avian) | 9.37 | 0.0210 |
| **ZBTB16** | zinc finger and BTB domain containing 16 | 9.34 | 0.0210 |
| **KCNS1** | potassium voltage-gated channel, delayed-rectifier, subfamily S, member 1 | 9.28 | 0.0214 |
| **PLXDC1** | plexin domain containing 1 | 9.24 | 0.0216 |
| **RAD54B** | RAD54 homolog B (S. cerevisiae) | 9.23 | 0.0216 |
| **NAPEPLD** | N-acyl phosphatidylethanolamine phospholipase D | 9.18 | 0.0219 |
| **HTR2C** | 5-hydroxytryptamine (serotonin) receptor 2C, G protein-coupled | -9.12 | 0.0220 |
| **NEFH** | neurofilament, heavy polypeptide | 9.12 | 0.0220 |
| **BHMT2** | betaine--homocysteine S-methyltransferase 2 | -9.02 | 0.0222 |
| **ECM1** | extracellular matrix protein 1 | 9.05 | 0.0222 |
| **PHYH** | phytanoyl-CoA 2-hydroxylase | 9.03 | 0.0222 |
| **CCDC8** | coiled-coil domain containing 8 | -8.88 | 0.0223 |
| **CTNNAL1** | catenin (cadherin-associated protein), alpha-like 1 | 8.87 | 0.0223 |
| **EPCAM** | epithelial cell adhesion molecule | -8.84 | 0.0224 |
| **ANKH** | ankylosis, progressive homolog (mouse) | 8.79 | 0.0225 |
| **LIX1** | Lix1 homolog (chicken) | -8.80 | 0.0225 |
| **ROBO1** | roundabout, axon guidance receptor, homolog 1 (Drosophila) | -8.82 | 0.0225 |

| | | | |
|---|---|---|---|
| **TMEM132E** | transmembrane protein 132E | 8.79 | 0.0225 |
| **ST3GAL6** | ST3 beta-galactoside alpha-2,3-sialyltransferase 6 | 8.75 | 0.0225 |
| **FNDC4** | fibronectin type III domain containing 4 | 8.71 | 0.0229 |
| **CAMK2G** | calcium/calmodulin-dependent protein kinase II gamma | 8.71 | 0.0229 |
| **NIPAL2** | NIPA-like domain containing 2 | 8.69 | 0.0229 |
| **MR1** | major histocompatibility complex, class I-related | -8.61 | 0.0231 |
| **COCH** | coagulation factor C homolog, cochlin (Limulus polyphemus) | -8.54 | 0.0232 |
| **KIF2C** | kinesin family member 2C | 8.55 | 0.0232 |
| **PPARGC1A** | peroxisome proliferator-activated receptor gamma, coactivator 1 alpha | 8.55 | 0.0232 |
| **STEAP2** | STEAP family member 2, metalloreductase | 8.52 | 0.0234 |
| **TIMP3** | TIMP metallopeptidase inhibitor 3 | -8.46 | 0.0239 |
| **EFNA5** | ephrin-A5 | 8.44 | 0.0239 |
| **RIMKLA** | ribosomal modification protein rimK-like family member A | 8.44 | 0.0239 |
| **CABP1** | calcium binding protein 1 | 8.38 | 0.0243 |
| **SOHLH1** | spermatogenesis and oogenesis specific basic helix-loop-helix 1 | 8.37 | 0.0244 |
| **RRP7A** | ribosomal RNA processing 7 homolog A (S. cerevisiae) | -8.33 | 0.0247 |
| **NCK2** | NCK adaptor protein 2 | 8.26 | 0.0251 |
| **DMKN** | dermokine | 8.21 | 0.0253 |
| **MUCL1** | mucin-like 1 | 8.20 | 0.0253 |
| **P2RY1** | purinergic receptor P2Y, G-protein coupled, 1 | -8.20 | 0.0253 |
| **CHRNA2** | cholinergic receptor, nicotinic, alpha 2 (neuronal) | 8.07 | 0.0265 |
| **SIRT4** | sirtuin 4 | 8.06 | 0.0265 |
| **DNAJC12** | DnaJ (Hsp40) homolog, subfamily C, member 12 | -7.96 | 0.0269 |
| **ICA1** | islet cell autoantigen 1, 69kDa | 7.98 | 0.0269 |
| **LYPLAL1** | lysophospholipase-like 1 | -8.00 | 0.0269 |
| **PON3** | paraoxonase 3 | -7.96 | 0.0269 |
| **HR** | hairless homolog (mouse) | 7.95 | 0.0269 |
| **RCAN2** | regulator of calcineurin 2 | 7.91 | 0.0272 |
| **CASQ1** | calsequestrin 1 (fast-twitch, skeletal muscle) | 7.78 | 0.0274 |
| **CHGA** | chromogranin A (parathyroid secretory protein 1) | 7.85 | 0.0274 |
| **FSTL1** | follistatin-like 1 | 7.74 | 0.0274 |
| **GABRB2** | gamma-aminobutyric acid (GABA) A receptor, beta 2 | 7.75 | 0.0274 |
| **GLUD2** | glutamate dehydrogenase 2 | -7.83 | 0.0274 |
| **GNG10** | guanine nucleotide binding protein (G protein), gamma 10 | -7.80 | 0.0274 |
| **KCNE4** | potassium voltage-gated channel, Isk-related family, member 4 | -7.76 | 0.0274 |
| **PGRMC1** | progesterone receptor membrane component 1 | -7.79 | 0.0274 |
| **PTPN4** | protein tyrosine phosphatase, non-receptor type 4 (megakaryocyte) | 7.86 | 0.0274 |
| **SEMA7A** | semaphorin 7A, GPI membrane anchor (John Milton Hagen blood group) | 7.80 | 0.0274 |
| **PALLD** | palladin, cytoskeletal associated protein | -7.73 | 0.0274 |
| **ESRRG** | estrogen-related receptor gamma | 7.70 | 0.0276 |
| **GTF2F2** | general transcription factor IIF, polypeptide 2, 30kDa | -7.68 | 0.0278 |
| **N4BP2L1** | NEDD4 binding protein 2-like 1 | 7.67 | 0.0278 |
| **ATP4A** | ATPase, H+/K+ exchanging, alpha polypeptide | 7.64 | 0.0279 |
| **ASB2** | ankyrin repeat and SOCS box containing 2 | 7.63 | 0.0280 |
| **FGD4** | FYVE, RhoGEF and PH domain containing 4 | -7.62 | 0.0280 |
| **HCN1** | hyperpolarization activated cyclic nucleotide-gated potassium channel 1 | 7.62 | 0.0280 |
| **KLF9** | Kruppel-like factor 9 | 7.61 | 0.0280 |
| **SLC29A1** | solute carrier family 29 (nucleoside transporters), member 1 | 7.63 | 0.0280 |
| **DKK1** | dickkopf 1 homolog (Xenopus laevis) | 7.58 | 0.0283 |
| **GLRX** | glutaredoxin (thioltransferase) | 7.55 | 0.0286 |
| **ANKRD34C** | ankyrin repeat domain 34C | 7.48 | 0.0290 |
| **C12orf45** | chromosome 12 open reading frame 45 | -7.44 | 0.0290 |

| | | | |
|---|---|---|---|
| **RRAGB** | Ras-related GTP binding B | -7.42 | 0.0293 |
| **STS** | steroid sulfatase (microsomal), isozyme S | 7.41 | 0.0293 |
| **ADPRHL1** | ADP-ribosylhydrolase like 1 | 7.35 | 0.0295 |
| **IMPACT** | Impact homolog (mouse) | -7.36 | 0.0295 |
| **TMEM120A** | transmembrane protein 120A | -7.35 | 0.0295 |
| **WDFY4** | WDFY family member 4 | -7.35 | 0.0295 |
| **ZNF385D** | zinc finger protein 385D | 7.35 | 0.0295 |
| **GNAS** | GNAS complex locus | 7.33 | 0.0297 |
| **FBXO9** | F-box protein 9 | 7.31 | 0.0299 |
| **CPNE6** | copine VI (neuronal) | -7.25 | 0.0300 |
| **DNAJA4** | DnaJ (Hsp40) homolog, subfamily A, member 4 | -7.21 | 0.0300 |
| **FBXO33** | F-box protein 33 | 7.22 | 0.0300 |
| **IQCA1** | IQ motif containing with AAA domain 1 | -7.26 | 0.0300 |
| **LGI2** | leucine-rich repeat LGI family, member 2 | 7.22 | 0.0300 |
| **LRRC36** | leucine rich repeat containing 36 | -7.20 | 0.0300 |
| **MADCAM1** | mucosal vascular addressin cell adhesion molecule 1 | 7.24 | 0.0300 |
| **MGP** | matrix Gla protein | 7.28 | 0.0300 |
| **OR2L8** | olfactory receptor, family 2, subfamily L, member 8 | 7.23 | 0.0300 |
| **PHLDA2** | pleckstrin homology-like domain, family A, member 2 | 7.17 | 0.0300 |
| **RARB** | retinoic acid receptor, beta | 7.26 | 0.0300 |
| **RPP25** | ribonuclease P/MRP 25kDa subunit | 7.24 | 0.0300 |
| **STAC2** | SH3 and cysteine rich domain 2 | 7.25 | 0.0300 |
| **TRIM37** | tripartite motif containing 37 | 7.19 | 0.0300 |
| **WDR66** | WD repeat domain 66 | -7.26 | 0.0300 |
| **LOC642852** | uncharacterized LOC642852 | -7.15 | 0.0301 |
| **MAGI3** | membrane associated guanylate kinase, WW and PDZ domain containing 3 | 7.15 | 0.0301 |
| **SULT1C4** | sulfotransferase family, cytosolic, 1C, member 4 | -7.08 | 0.0304 |
| **PGAP1** | post-GPI attachment to proteins 1 | -7.05 | 0.0306 |
| **PTGS1** | prostaglandin-endoperoxide synthase 1 (prostaglandin G/H synthase and cyclooxygen | 7.04 | 0.0306 |
| **CPE** | carboxypeptidase E | -7.00 | 0.0310 |
| **TSPAN5** | tetraspanin 5 | 6.99 | 0.0310 |
| **POU6F2** | POU class 6 homeobox 2 | 6.96 | 0.0311 |
| **SCN8A** | sodium channel, voltage gated, type VIII, alpha subunit | 6.97 | 0.0311 |
| **SORL1** | sortilin-related receptor, L(DLR class) A repeats containing | 6.95 | 0.0313 |
| **ENTPD4** | ectonucleoside triphosphate diphosphohydrolase 4 | 6.94 | 0.0314 |
| **MINK1** | misshapen-like kinase 1 | -6.92 | 0.0315 |
| **PARD3B** | par-3 partitioning defective 3 homolog B (C. elegans) | -6.89 | 0.0317 |
| **RNF150** | ring finger protein 150 | -6.90 | 0.0317 |
| **TMEM176A** | transmembrane protein 176A | -6.90 | 0.0317 |
| **ARL4C** | ADP-ribosylation factor-like 4C | 6.87 | 0.0317 |
| **KCNJ11** | potassium inwardly-rectifying channel, subfamily J, member 11 | 6.86 | 0.0317 |
| **PREX2** | phosphatidylinositol-3,4,5-trisphosphate-dependent Rac exchange factor 2 | -6.86 | 0.0317 |
| **LGALS1** | lectin, galactoside-binding, soluble, 1 | 6.85 | 0.0318 |
| **RHOC** | ras homolog family member C | -6.84 | 0.0319 |
| **RYR3** | ryanodine receptor 3 | -6.81 | 0.0322 |
| **DQX1** | DEAQ box RNA-dependent ATPase 1 | 6.79 | 0.0323 |
| **MYH7B** | myosin, heavy chain 7B, cardiac muscle, beta | 6.79 | 0.0323 |
| **IGFBP2** | insulin-like growth factor binding protein 2, 36kDa | 6.78 | 0.0324 |
| **SYT12** | synaptotagmin XII | 6.75 | 0.0325 |
| **TPBG** | trophoblast glycoprotein | 6.76 | 0.0325 |
| **LRRN1** | leucine rich repeat neuronal 1 | -6.73 | 0.0330 |
| **STK17A** | serine/threonine kinase 17a | -6.72 | 0.0330 |

| | | | |
|---|---|---|---|
| **UBE2QL1** | ubiquitin-conjugating enzyme E2Q family-like 1 | 6.71 | 0.0330 |
| **NT5M** | 5,3-nucleotidase, mitochondrial | 6.69 | 0.0332 |
| **PCP4L1** | Purkinje cell protein 4 like 1 | 6.65 | 0.0333 |
| **MKX** | mohawk homeobox | 6.65 | 0.0333 |
| **SLC16A2** | solute carrier family 16, member 2 (thyroid hormone transporter) | -6.64 | 0.0333 |
| **MT1F** | metallothionein 1F | -6.62 | 0.0336 |
| **PTPN3** | protein tyrosine phosphatase, non-receptor type 3 | 6.59 | 0.0336 |
| **POSTN** | periostin, osteoblast specific factor | 6.56 | 0.0338 |
| **UG0898H09** | uncharacterized LOC643763 | -6.56 | 0.0338 |
| **ADCYAP1R1** | adenylate cyclase activating polypeptide 1 (pituitary) receptor type I | -6.55 | 0.0340 |
| **ECHDC3** | enoyl CoA hydratase domain containing 3 | -6.54 | 0.0340 |
| **FABP5P3** | fatty acid binding protein 5 pseudogene 3 | -6.53 | 0.0342 |
| **SLC25A5** | solute carrier family 25 (mitochondrial carrier; adenine nucleotide translocator), mem | 6.53 | 0.0342 |
| **CDS1** | CDP-diacylglycerol synthase (phosphatidate cytidylyltransferase) 1 | 6.51 | 0.0343 |
| **APOC1** | apolipoprotein C-I | -6.50 | 0.0343 |
| **BTN3A2** | butyrophilin, subfamily 3, member A2 | -6.50 | 0.0343 |
| **CTSO** | cathepsin O | -6.50 | 0.0343 |
| **RET** | ret proto-oncogene | 6.48 | 0.0345 |
| **SMPX** | small muscle protein, X-linked | 6.47 | 0.0346 |
| **TUBGCP5** | tubulin, gamma complex associated protein 5 | 6.47 | 0.0346 |
| **FGF9** | fibroblast growth factor 9 (glia-activating factor) | 6.45 | 0.0348 |
| **PRDM2** | PR domain containing 2, with ZNF domain | 6.44 | 0.0348 |
| **SERTAD4** | SERTA domain containing 4 | 6.42 | 0.0348 |
| **TMEM136** | transmembrane protein 136 | -6.42 | 0.0348 |
| **LYPD5** | LY6/PLAUR domain containing 5 | 6.39 | 0.0350 |
| **ARL9** | ADP-ribosylation factor-like 9 | 6.35 | 0.0353 |
| **CXorf57** | chromosome X open reading frame 57 | -6.35 | 0.0353 |
| **PTH2R** | parathyroid hormone 2 receptor | 6.35 | 0.0353 |
| **P2RX6P** | purinergic receptor P2X, ligand-gated ion channel, 6 pseudogene | 6.35 | 0.0353 |
| **PKIA** | protein kinase (cAMP-dependent, catalytic) inhibitor alpha | -6.33 | 0.0353 |
| **SLC16A7** | solute carrier family 16, member 7 (monocarboxylic acid transporter 2) | 6.34 | 0.0353 |
| **TNNT2** | troponin T type 2 (cardiac) | 6.33 | 0.0353 |
| **KCTD10** | potassium channel tetramerisation domain containing 10 | -6.32 | 0.0354 |
| **CREM** | cAMP responsive element modulator | -6.28 | 0.0355 |
| **EDNRB** | endothelin receptor type B | -6.28 | 0.0355 |
| **IFFO1** | intermediate filament family orphan 1 | 6.28 | 0.0355 |
| **IRS1** | insulin receptor substrate 1 | 6.31 | 0.0355 |
| **NECAB3** | N-terminal EF-hand calcium binding protein 3 | 6.26 | 0.0355 |
| **PEA15** | phosphoprotein enriched in astrocytes 15 | -6.27 | 0.0355 |
| **RSPO3** | R-spondin 3 | -6.29 | 0.0355 |
| **TFEC** | transcription factor EC | -6.26 | 0.0356 |
| **EPHX1** | epoxide hydrolase 1, microsomal (xenobiotic) | -6.24 | 0.0357 |
| **GSDMB** | gasdermin B | 6.24 | 0.0357 |
| **OSTF1** | osteoclast stimulating factor 1 | 6.24 | 0.0357 |
| **HTR1E** | 5-hydroxytryptamine (serotonin) receptor 1E, G protein-coupled | 6.22 | 0.0358 |
| **FZD1** | frizzled family receptor 1 | -6.20 | 0.0360 |
| **THEMIS** | thymocyte selection associated | 6.18 | 0.0363 |
| **ALDH1A3** | aldehyde dehydrogenase 1 family, member A3 | 6.17 | 0.0364 |
| **SPHKAP** | SPHK1 interactor, AKAP domain containing | -6.16 | 0.0365 |
| **RHOBTB2** | Rho-related BTB domain containing 2 | 6.12 | 0.0372 |
| **RTKN2** | rhotekin 2 | 6.11 | 0.0372 |
| **DLG5** | discs, large homolog 5 (Drosophila) | 6.10 | 0.0372 |

| | | | |
|---|---|---|---|
| **NIF3L1** | NIF3 NGG1 interacting factor 3-like 1 (S. cerevisiae) | 6.10 | 0.0373 |
| **E2F5** | E2F transcription factor 5, p130-binding | -6.09 | 0.0373 |
| **KLHL8** | kelch-like 8 (Drosophila) | 6.08 | 0.0373 |
| **TTR** | transthyretin | -6.09 | 0.0373 |
| **PELI3** | pellino E3 ubiquitin protein ligase family member 3 | 6.06 | 0.0375 |
| **ST8SIA1** | ST8 alpha-N-acetyl-neuraminide alpha-2,8-sialyltransferase 1 | 6.06 | 0.0375 |
| **PNMT** | phenylethanolamine N-methyltransferase | -6.04 | 0.0376 |
| **ZMAT1** | zinc finger, matrin-type 1 | -6.03 | 0.0377 |
| **FNBP1L** | formin binding protein 1-like | -5.99 | 0.0381 |
| **GPR161** | G protein-coupled receptor 161 | 5.98 | 0.0382 |
| **SH3BGRL3** | SH3 domain binding glutamic acid-rich protein like 3 | -5.97 | 0.0382 |
| **CMYA5** | cardiomyopathy associated 5 | 5.97 | 0.0382 |
| **FABP6** | fatty acid binding protein 6, ileal | -5.96 | 0.0384 |
| **CCDC85A** | coiled-coil domain containing 85A | 5.95 | 0.0385 |
| **LUZP1** | leucine zipper protein 1 | 5.94 | 0.0386 |
| **ANKRD50** | ankyrin repeat domain 50 | -5.91 | 0.0390 |
| **CADM1** | cell adhesion molecule 1 | -5.91 | 0.0391 |
| **MILR1** | mast cell immunoglobulin-like receptor 1 | -5.91 | 0.0391 |
| **DCN** | decorin | -5.90 | 0.0392 |
| **DAPL1** | death associated protein-like 1 | -5.87 | 0.0396 |
| **RHBDL3** | rhomboid, veinlet-like 3 (Drosophila) | 5.86 | 0.0396 |
| **RAB3C** | RAB3C, member RAS oncogene family | -5.86 | 0.0398 |
| **LRCH2** | leucine-rich repeats and calponin homology (CH) domain containing 2 | -5.85 | 0.0399 |
| **SKAP1** | src kinase associated phosphoprotein 1 | -5.83 | 0.0401 |
| **PLEKHM2** | pleckstrin homology domain containing, family M (with RUN domain) member 2 | 5.82 | 0.0402 |
| **ESYT1** | extended synaptotagmin-like protein 1 | 5.81 | 0.0405 |
| **CPNE3** | copine III | -5.80 | 0.0406 |
| **FAM196A** | family with sequence similarity 196, member A | -5.80 | 0.0406 |
| **LY86-AS1** | LY86 antisense RNA 1 (non-protein coding) | 5.78 | 0.0407 |
| **ACAN** | aggrecan | 5.75 | 0.0411 |
| **SLA** | Src-like-adaptor | -5.73 | 0.0413 |
| **SLCO4A1** | solute carrier organic anion transporter family, member 4A1 | 5.72 | 0.0415 |
| **PDIA5** | protein disulfide isomerase family A, member 5 | 5.67 | 0.0424 |
| **INTU** | inturned planar cell polarity effector homolog (Drosophila) | -5.65 | 0.0427 |
| **ANKRD6** | ankyrin repeat domain 6 | -5.65 | 0.0427 |
| **AQP9** | aquaporin 9 | 5.64 | 0.0428 |
| **LOXL1** | lysyl oxidase-like 1 | -5.63 | 0.0428 |
| **CMPK1** | cytidine monophosphate (UMP-CMP) kinase 1, cytosolic | -5.62 | 0.0429 |
| **DIRAS3** | DIRAS family, GTP-binding RAS-like 3 | -5.61 | 0.0429 |
| **SESN3** | sestrin 3 | -5.61 | 0.0429 |
| **SYTL2** | synaptotagmin-like 2 | 5.62 | 0.0429 |
| **FAM43A** | family with sequence similarity 43, member A | 5.58 | 0.0431 |
| **OPHN1** | oligophrenin 1 | -5.57 | 0.0431 |
| **SLC22A9** | solute carrier family 22 (organic anion transporter), member 9 | 5.57 | 0.0431 |
| **SULF2** | sulfatase 2 | -5.57 | 0.0431 |
| **DPP8** | dipeptidyl-peptidase 8 | 5.50 | 0.0439 |
| **KIAA1107** | KIAA1107 | 5.51 | 0.0439 |
| **SLC25A12** | solute carrier family 25 (aspartate/glutamate carrier), member 12 | 5.49 | 0.0441 |
| **DTWD1** | DTW domain containing 1 | -5.48 | 0.0443 |
| **NKAIN3** | Na+/K+ transporting ATPase interacting 3 | -5.48 | 0.0443 |
| **KNG1** | kininogen 1 | 5.46 | 0.0444 |
| **NAT8L** | N-acetyltransferase 8-like (GCN5-related, putative) | 5.46 | 0.0444 |

| | | | |
|---|---|---|---|
| **TSHZ1** | teashirt zinc finger homeobox 1 | 5.46 | 0.0444 |
| **CHID1** | chitinase domain containing 1 | -5.43 | 0.0448 |
| **FMN1** | formin 1 | 5.42 | 0.0448 |
| **GPR137C** | G protein-coupled receptor 137C | 5.42 | 0.0448 |
| **HES4** | hairy and enhancer of split 4 (Drosophila) | -5.43 | 0.0448 |
| **CRTAC1** | cartilage acidic protein 1 | 5.41 | 0.0449 |
| **NUPR1** | nuclear protein, transcriptional regulator, 1 | -5.40 | 0.0449 |
| **CD63** | CD63 molecule | -5.39 | 0.0450 |
| **KCNA5** | potassium voltage-gated channel, shaker-related subfamily, member 5 | 5.39 | 0.0451 |
| **STARD5** | StAR-related lipid transfer (START) domain containing 5 | 5.39 | 0.0451 |
| **FNDC5** | fibronectin type III domain containing 5 | 5.37 | 0.0452 |
| **ANK1** | ankyrin 1, erythrocytic | 5.36 | 0.0455 |
| **C17orf75** | chromosome 17 open reading frame 75 | 5.34 | 0.0459 |
| **CDC42EP3** | CDC42 effector protein (Rho GTPase binding) 3 | 5.32 | 0.0459 |
| **RILP** | Rab interacting lysosomal protein | 5.32 | 0.0459 |
| **ENKUR** | enkurin, TRPC channel interacting protein | -5.30 | 0.0461 |
| **FBLN2** | fibulin 2 | -5.30 | 0.0461 |
| **ABTB1** | ankyrin repeat and BTB (POZ) domain containing 1 | 5.29 | 0.0462 |
| **PYGL** | phosphorylase, glycogen, liver | -5.29 | 0.0462 |
| **FAM117A** | family with sequence similarity 117, member A | -5.28 | 0.0464 |
| **IL33** | interleukin 33 | -5.26 | 0.0466 |
| **SLC17A6** | solute carrier family 17 (sodium-dependent inorganic phosphate cotransporter), mem | 5.26 | 0.0466 |
| **ENO2** | enolase 2 (gamma, neuronal) | 5.25 | 0.0467 |
| **MET** | met proto-oncogene (hepatocyte growth factor receptor) | 5.23 | 0.0468 |
| **PRKCD** | protein kinase C, delta | -5.23 | 0.0468 |
| **BAIAP2L2** | BAI1-associated protein 2-like 2 | 5.22 | 0.0472 |
| **CHAF1A** | chromatin assembly factor 1, subunit A (p150) | 5.22 | 0.0472 |
| **SLC24A2** | solute carrier family 24 (sodium/potassium/calcium exchanger), member 2 | 5.20 | 0.0477 |
| **RXFP1** | relaxin/insulin-like family peptide receptor 1 | 5.19 | 0.0479 |
| **WDR1** | WD repeat domain 1 | -5.15 | 0.0484 |
| **CHRNA3** | cholinergic receptor, nicotinic, alpha 3 (neuronal) | -5.14 | 0.0486 |
| **GSTM2** | glutathione S-transferase mu 2 (muscle) | -5.14 | 0.0487 |
| **CACNB4** | calcium channel, voltage-dependent, beta 4 subunit | 5.11 | 0.0492 |
| **MOGAT1** | monoacylglycerol O-acyltransferase 1 | 5.11 | 0.0492 |
| **MAT2B** | methionine adenosyltransferase II, beta | 5.10 | 0.0493 |
| **KCNH7** | potassium voltage-gated channel, subfamily H (eag-related), member 7 | 5.08 | 0.0496 |
| **SETBP1** | SET binding protein 1 | 5.08 | 0.0497 |
| **FREM3** | FRAS1 related extracellular matrix 3 | 5.07 | 0.0498 |
| **VAV1** | vav 1 guanine nucleotide exchange factor | -5.07 | 0.0499 |
