## Supplementary Materials for "An expanding manifold in transmodal regions characterizes adolescent reconfiguration of structural connectome organization"

### Supporting Information

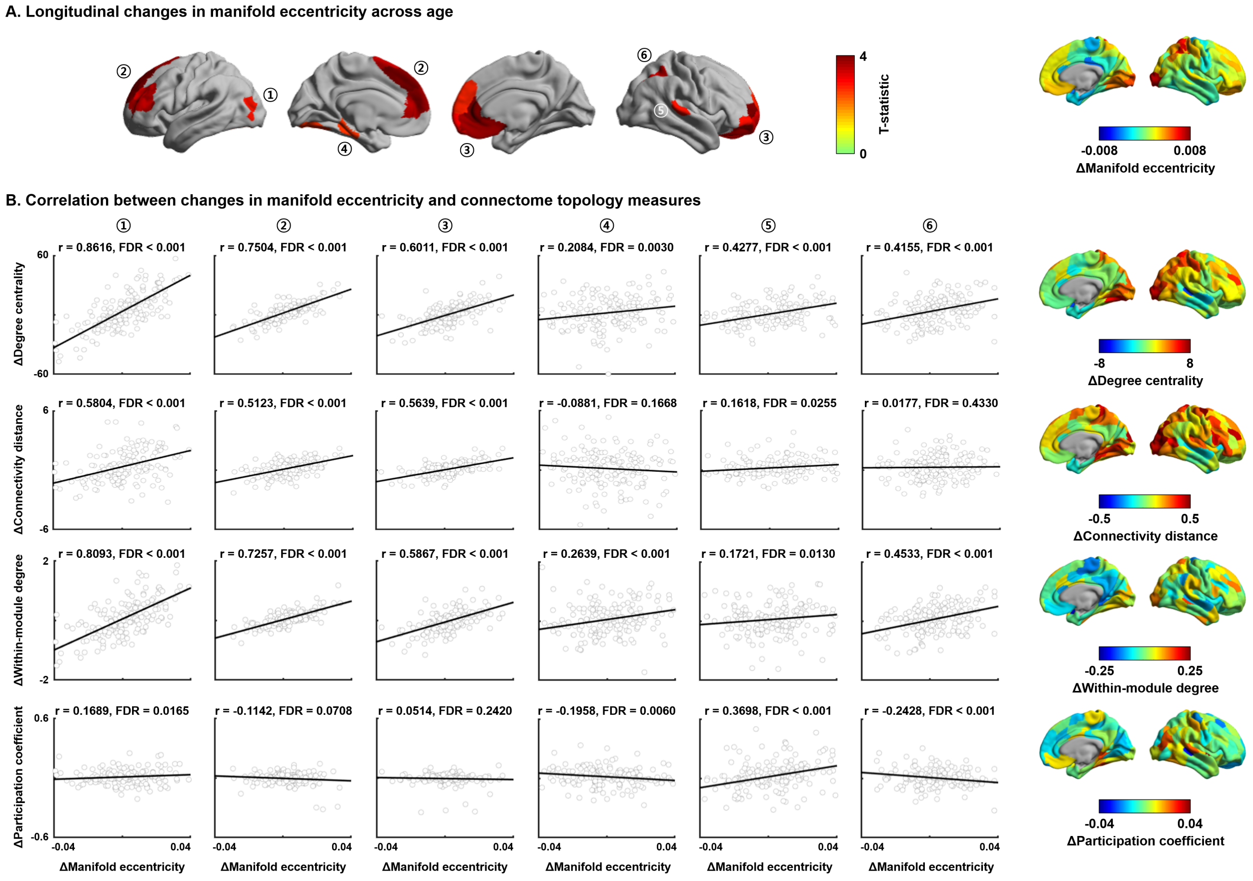


**Fig. S1 | Association between structural connectome manifold and connectome topology measures. (A)** Six clusters defined within the identified regions that showed significant age-related changes in manifold eccentricity (see *Fig. 1C*). **(B)** Associations between within-subject changes in manifold eccentricity and those of each connectome topology measure. Brain surfaces on the right side represent changes in each measure between baseline and follow-up. Significance was calculated using a false discovery rate (FDR).

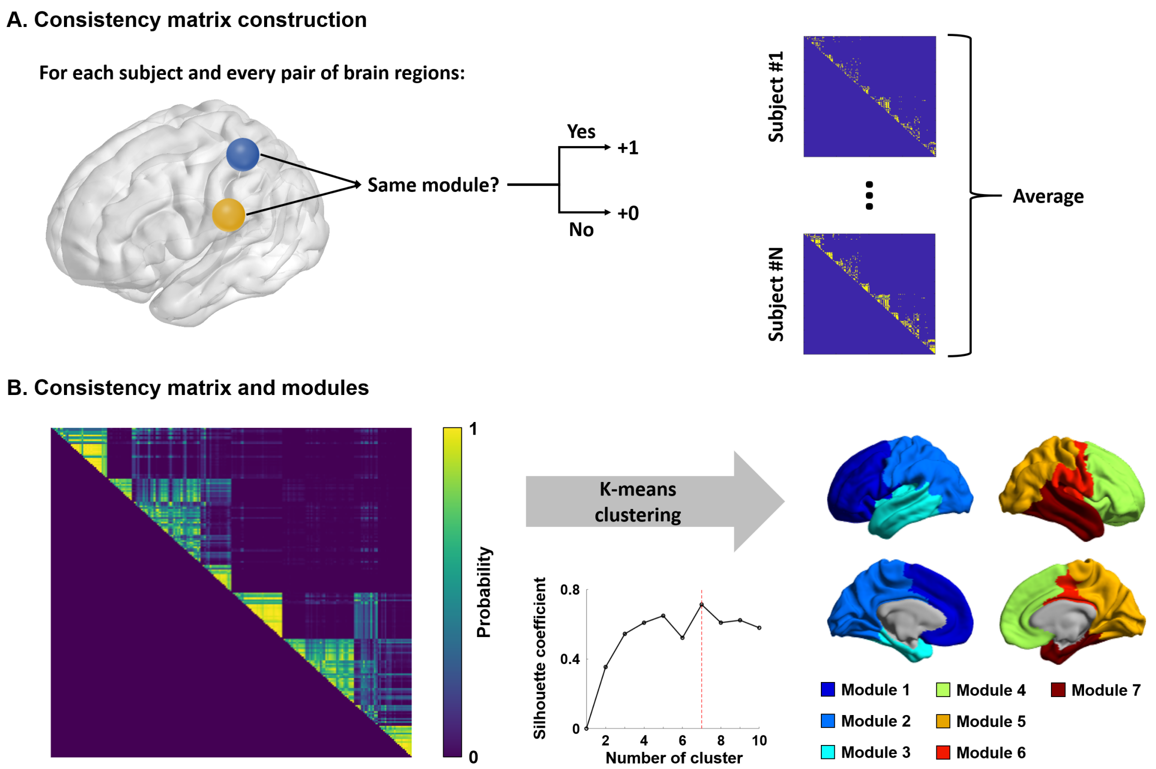


**Fig. S2 | Modular structures. (A)** Pipeline for constructing consistency matrix. We constructed individual subject-wise consistency matrix by considering whether two different nodes were involved in the same module. **(B)** Group-wise consistency matrix was constructed by averaging subject-wise consistency matrices. The k-means clustering with silhouette coefficient was used for defining modules. Seven modules on the brain surface are reported on the right side.

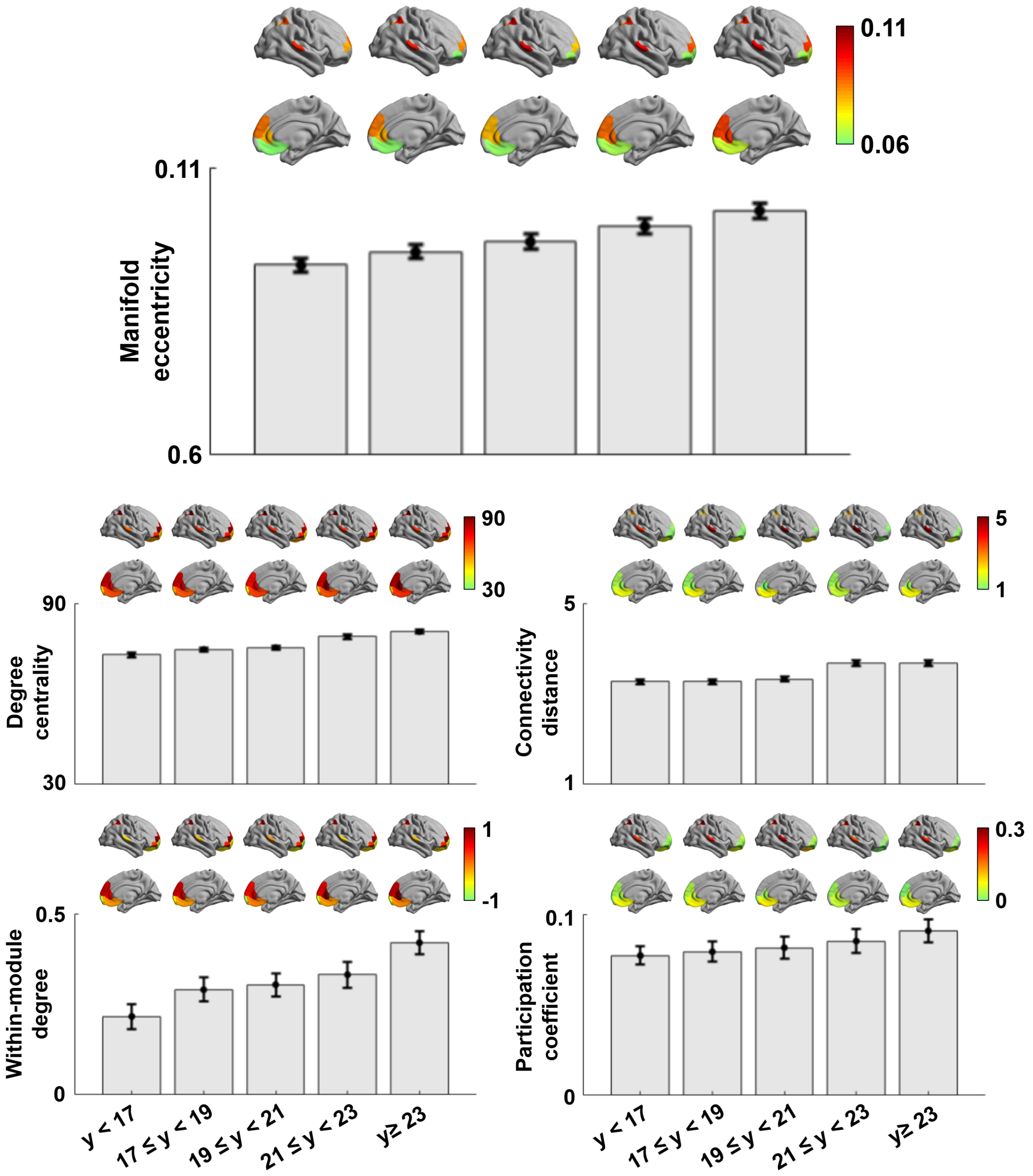


**Fig. S3 | Age-related trends in connectome topology measures.** Age-related changes in manifold eccentricity, degree centrality, connectivity distance, within-module degree, and participation coefficient. *Abbreviation:* y, years.

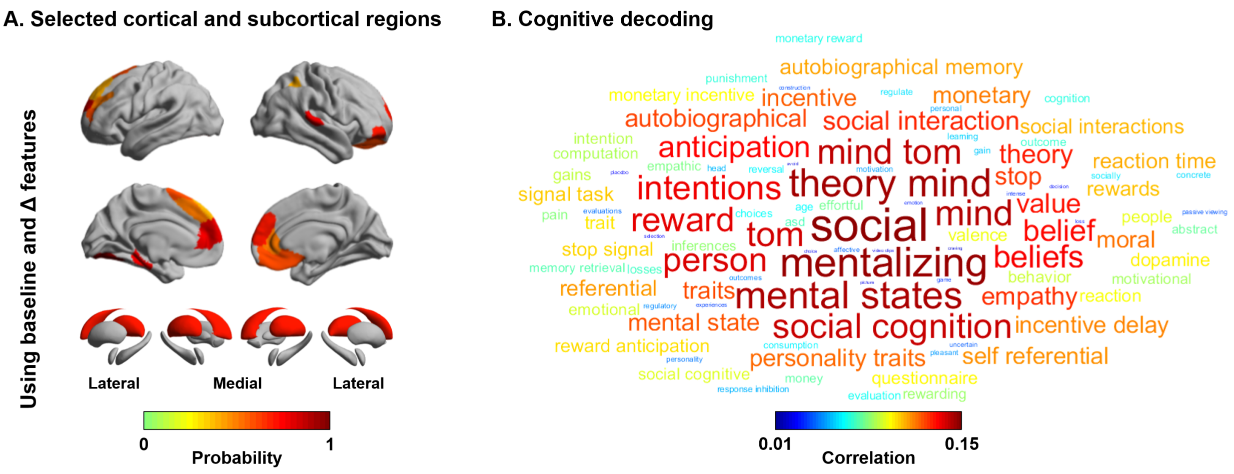


**Fig. S4 | Cognitive decoding of the selected regions for IQ prediction. (A)** Probability of selected cortical and subcortical regions for predicting future IQ using both baseline and maturational changes (see *Fig. 5*). **(B)** A word cloud derived by cognitive decoding using NeuroSynth (Yarkoni et al., 2011).

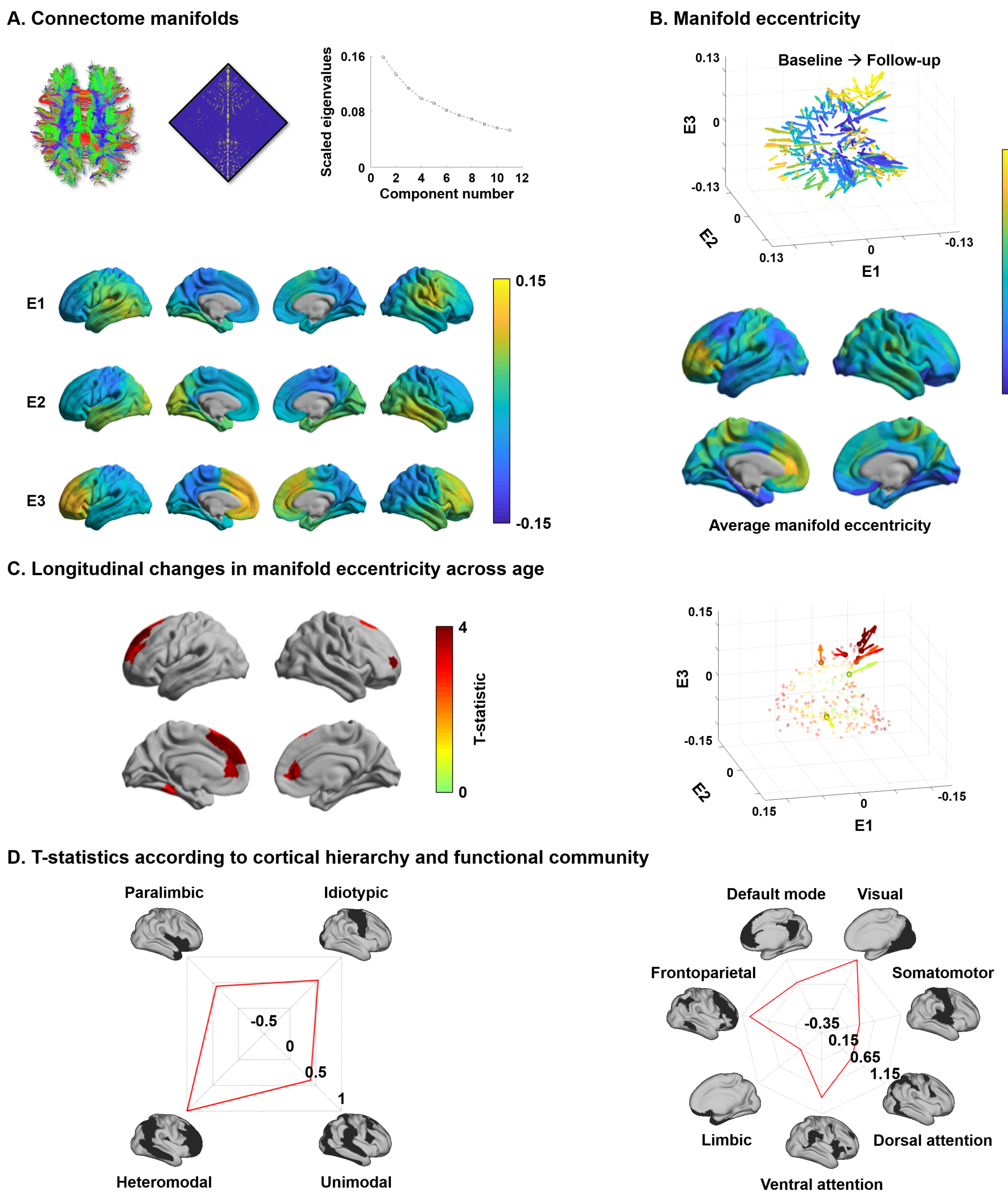


**Fig. S5 | Structural connectome manifolds using Schaefer 300 atlas. (A-D)** Main findings were replicated using a different parcellation scale. For details, see *Fig. 1*.

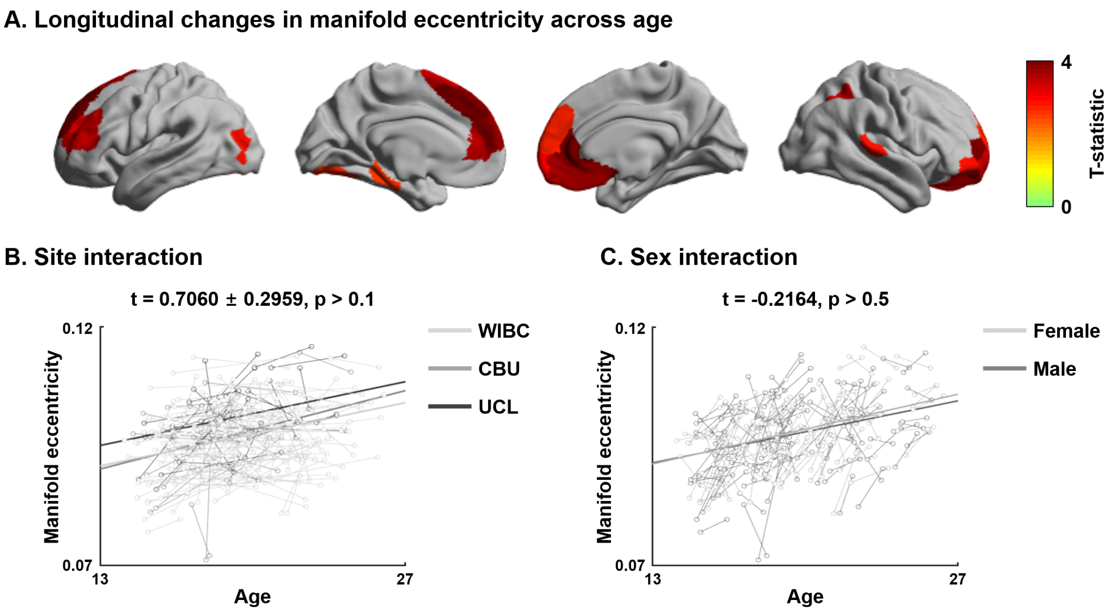


**Fig. S6 | Sensitivity analysis for site and sex. (A)** The t-statistics of identified regions that showed significant age-related changes in manifold eccentricity. **(B)** Interaction effects of the relationship between age and manifold eccentricity for sites and **(C)** biological sexes. *Abbreviations:* WIBC, Wolfson Brain Imaging Centre; CBU, MRC Cognition and Brain Sciences Unit; UCL, University College London.

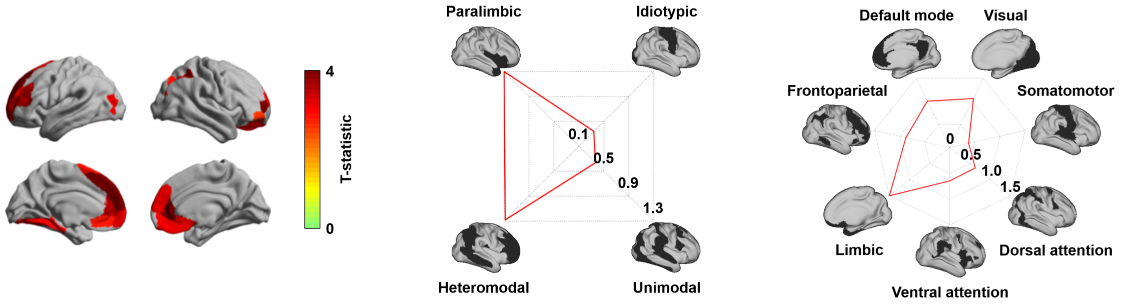


**Fig. S7 | Longitudinal changes in manifold eccentricity after excluding participants with the lowest correspondence to template manifolds.** The t-statistics of regions showing significant longitudinal changes in manifold eccentricity across age are reported on brain surface. Effects are stratified with respect to levels of cortical hierarchy (Mesulam, 1998) and intrinsic functional communities (Yeo et al., 2011).

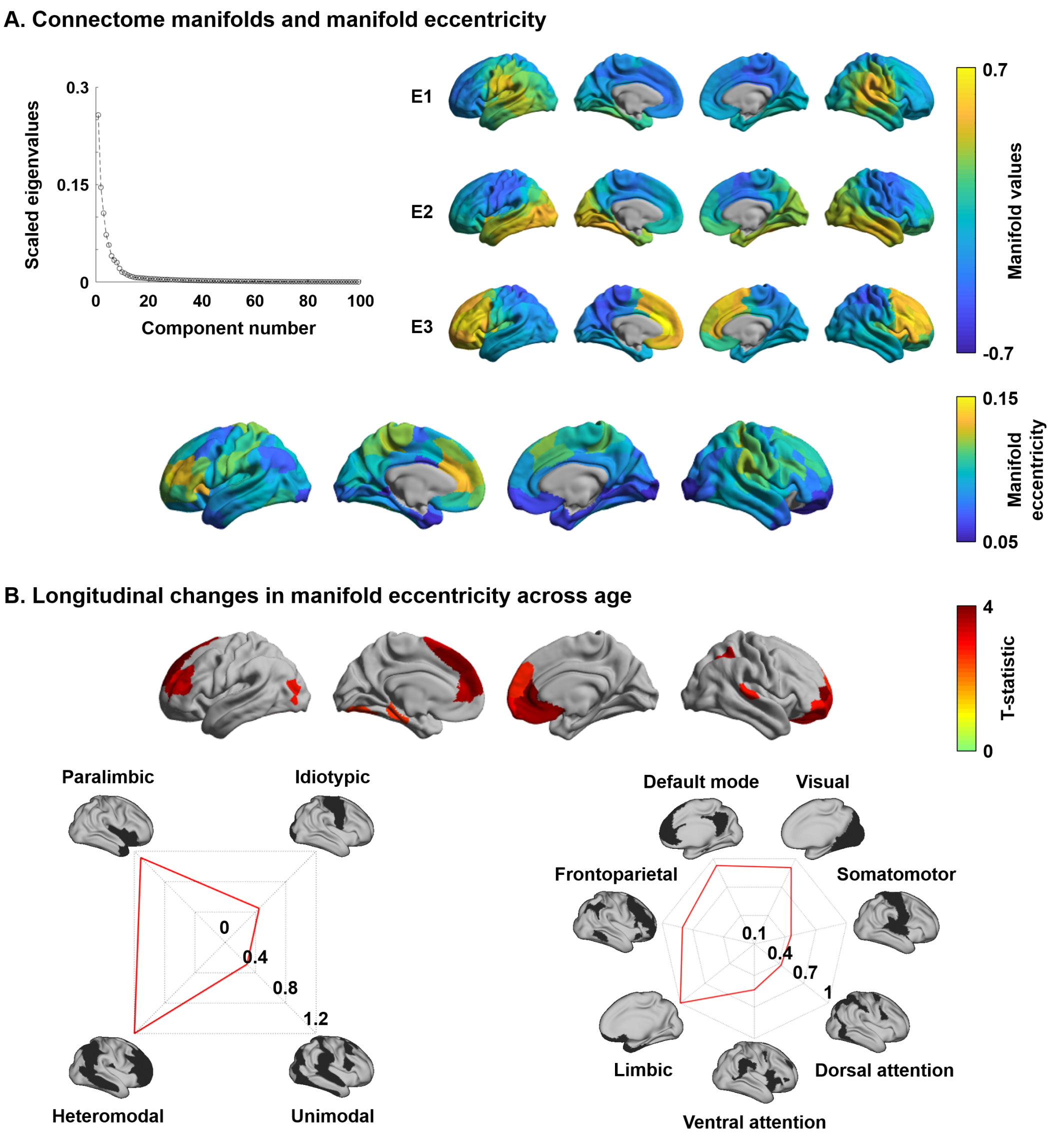


**Fig. S8 | Structural connectome manifolds generated using principal component analysis. (A)** Scree plot showing eigenvalue decay, and the first three eigenvectors (E1, E2, E3) are shown on brain surfaces. Manifold eccentricity is shown on the bottom. **(B)** Surface plots displaying t-statistics of regions showing significant longitudinal changes in manifold eccentricity across age. The effects are stratified with respect to levels of cortical hierarchy (Mesulam, 1998) and functional community (Yeo et al., 2011).

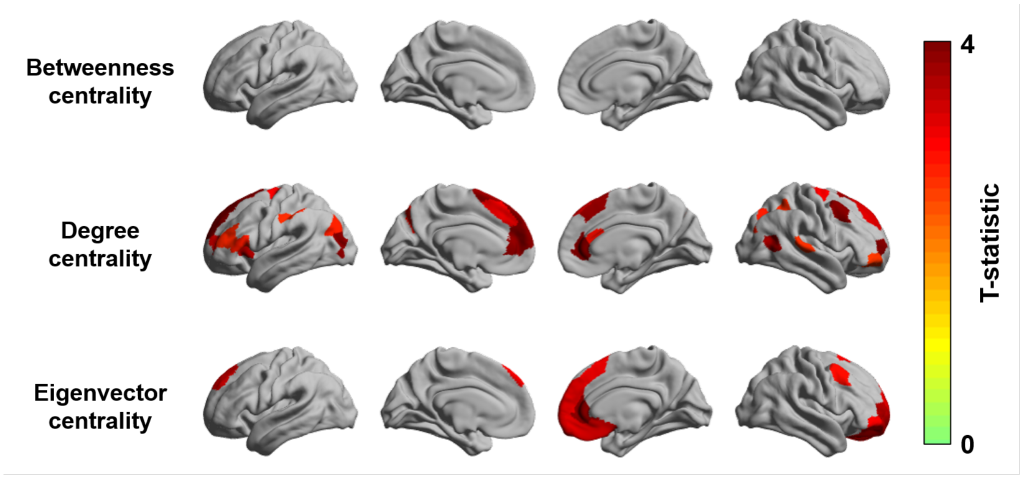


**Fig. S9 | Longitudinal changes in graph measures across age.** Cortical surface map showing t-statistics of regions showing significant age-related longitudinal changes in betweenness, degree, and eigenvector centrality.

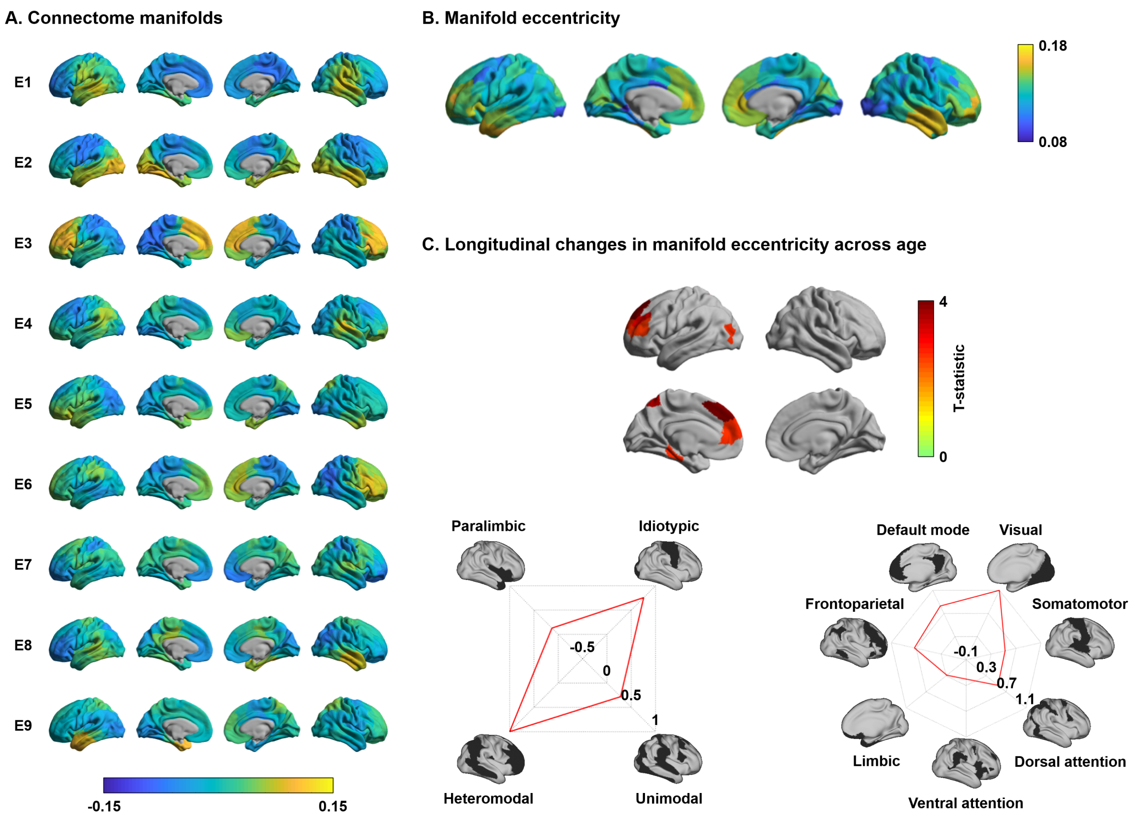


**Fig. S10 | Longitudinal changes in manifold eccentricity calculated using all eigenvectors. (A)** The generated eigenvectors (E1–E9) and **(B)** manifold eccentricity calculated using all of them. **(C)** The t-statistics of regions showing significant longitudinal changes in manifold eccentricity across age are reported on brain surface. The effects are stratified along cortical hierarchy (Mesulam, 1998) and functional community (Yeo et al., 2011).

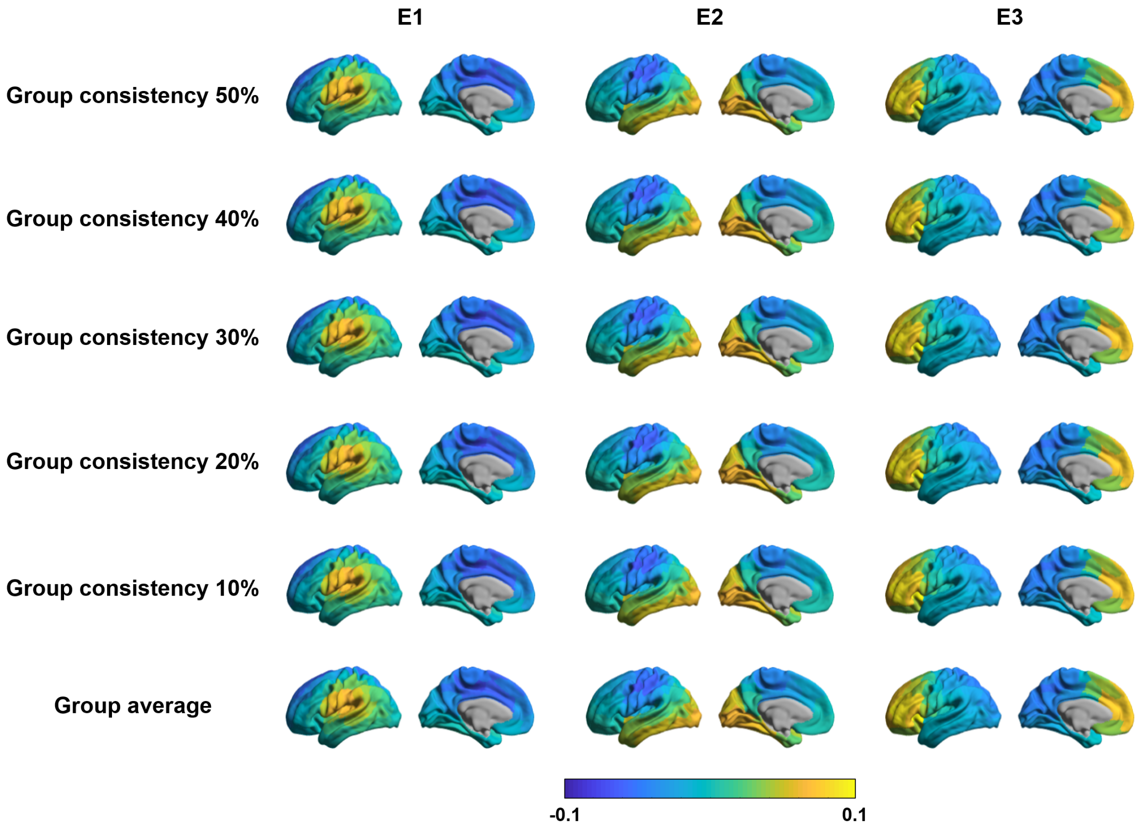


**Fig. S11 | Connectome manifolds estimated using group consistency method.** Spatial maps of three eigenvectors derived from group representative structural connectivity matrices based on different consistency thresholds are reported.

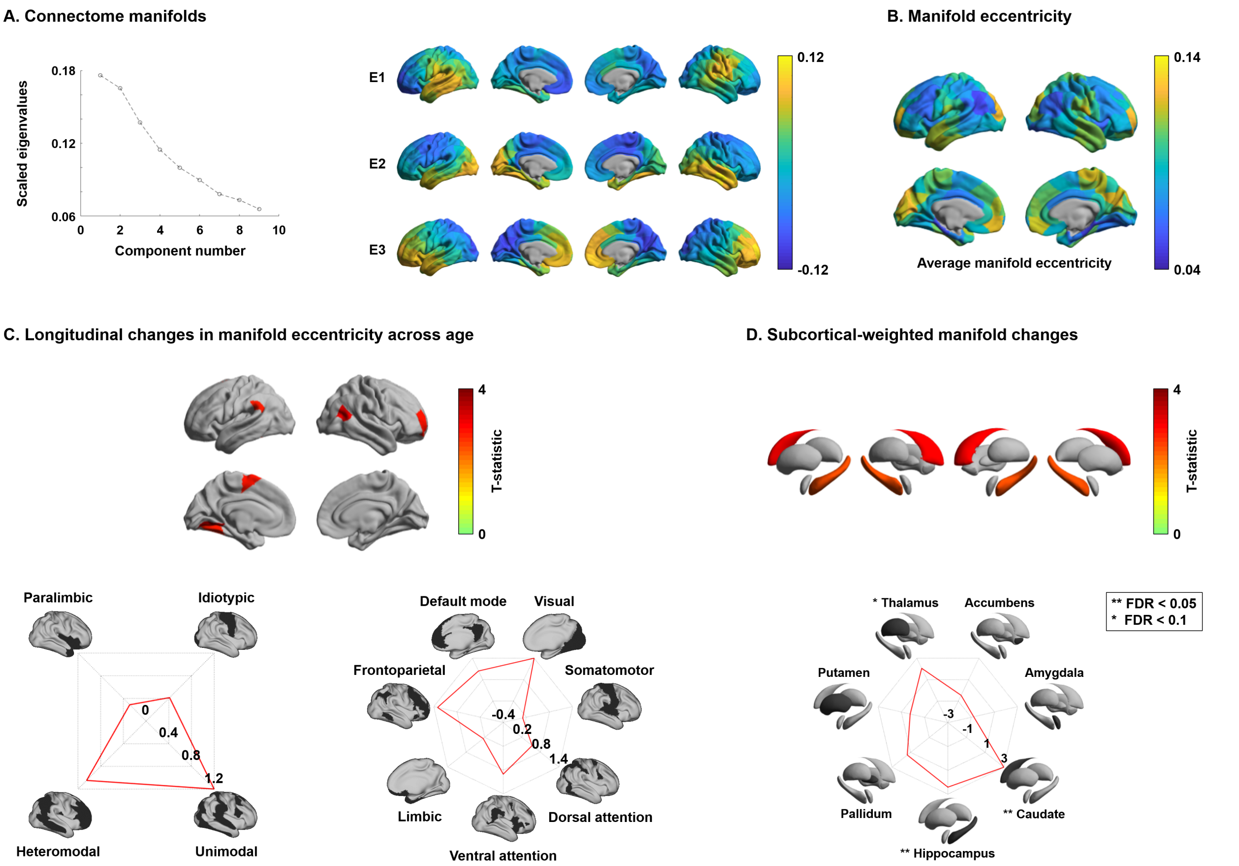


**Fig. S12 | Structural connectome manifolds using a structural parcellation. (A)** A scree plot shows eigenvalues of each component, and the first three eigenvectors (E1, E2, E3) are shown on brain surfaces. **(B)** Manifold eccentricity. **(C)** The t-statistics of regions showing significant longitudinal changes in manifold eccentricity and **(D)** subcortical-weighted manifolds across age. The effects of manifold eccentricity are stratified along cortical hierarchy (Mesulam, 1998) and functional community (Yeo et al., 2011). For details, see *Fig. 1*.

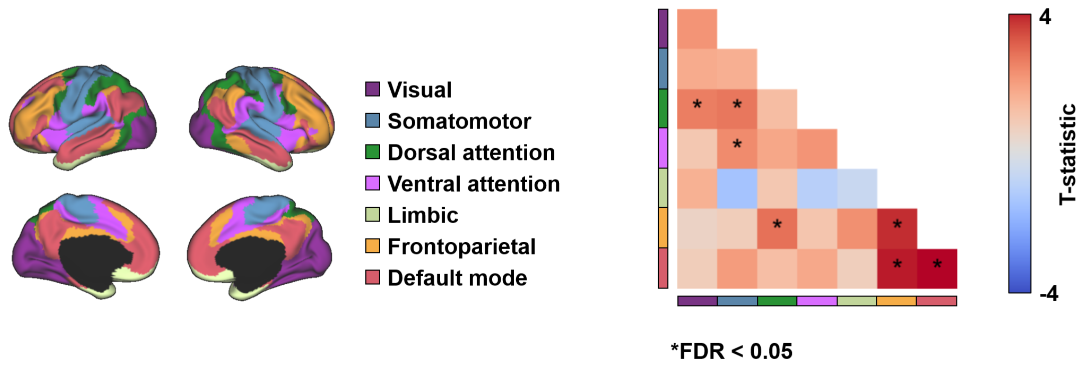


**Fig. S13 | Longitudinal changes in edge weights of structural connectome.** Findings were stratified relative to seven intrinsic functional communities (Yeo et al., 2011). The matrix displays t-statistics of connections showing longitudinal changes in edge weights, and significant (FDR < 0.05) results are marked with asterisks.

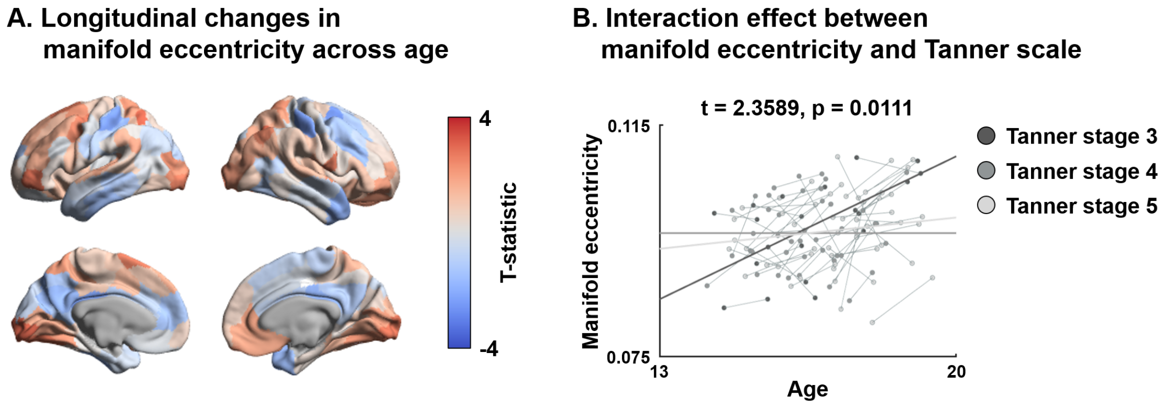


**Fig. S14 | Longitudinal changes in manifold eccentricity using a subset of participants who completed Tanner scale. (A)** Cortex t-statistics of age-related longitudinal effect. **(B)** Interaction effect between manifold eccentricity and Tanner scale. Colors of dots indicate Tanner stage of individuals, and lines indicate linear correlations between age and manifold eccentricity for individuals with the same Tanner stage.

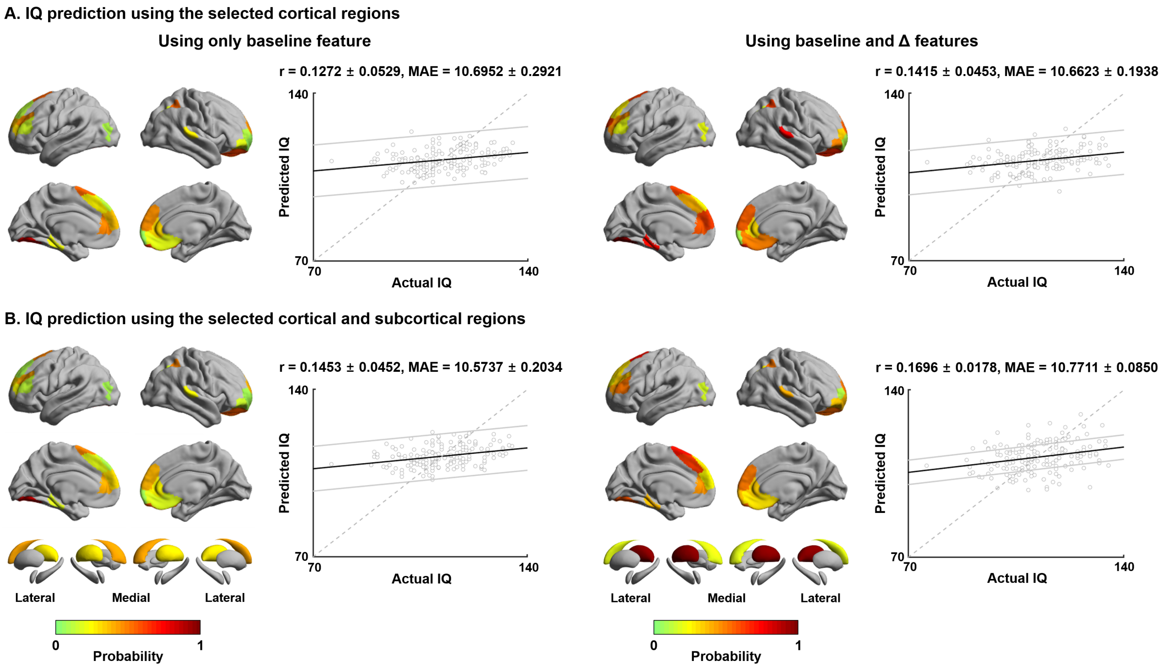


**Fig. S15 | IQ prediction using regression tree approach. (A)** The prediction performance using cortical features. Probability of selected brain regions across ten-fold cross-validation and 100 repetitions for predicting future IQ are represented on brain surfaces, and correlations between actual and predicted IQ are reported with scatter plots. **(B)** The prediction performance when both cortical and subcortical features were considered. For details, see *Fig. 5*.

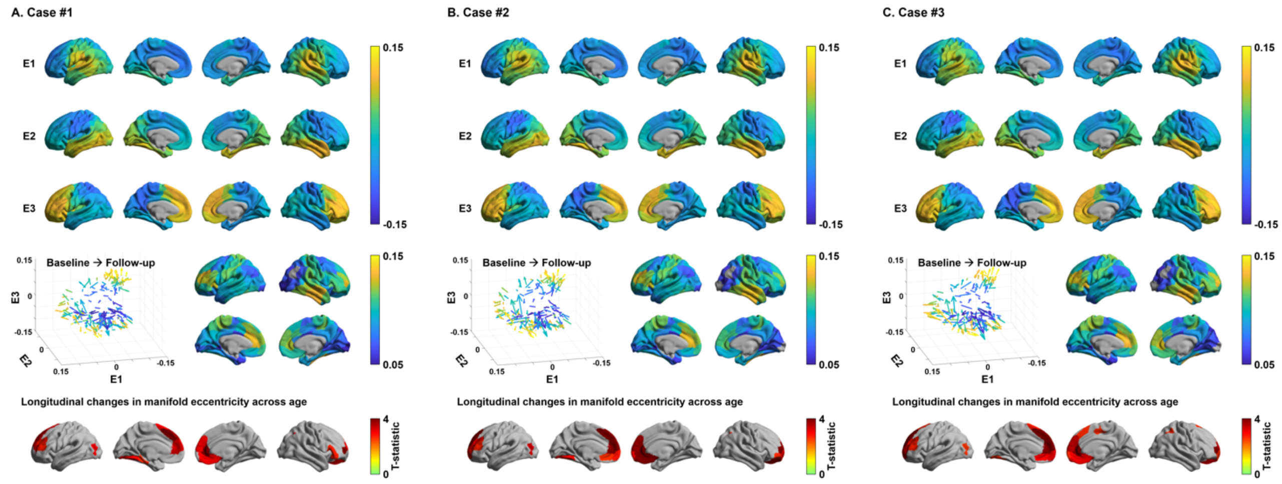


**Fig. S16 | Structural connectome manifolds using different template dataset. (A-C)** Structural connectome manifolds and association to age using different template cohort. Three representative cases are reported. For details, see *Figure 1*.

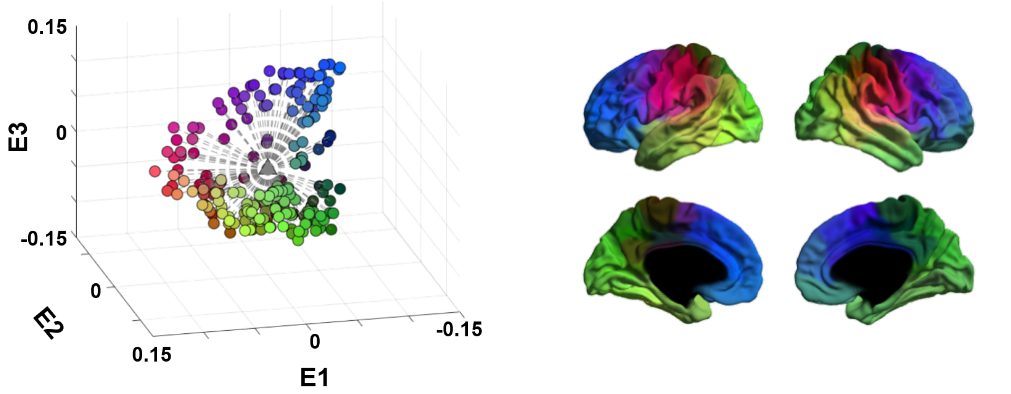


**Fig. S17 | A schema of manifold eccentricity for three eigenvectors.** Each dot in the scatter plot represents a single brain region, and the colors matched with regions on the brain surface. The triangle in the middle of the scatter plot is the manifold origin, and all brain regions (*i.e.,* dots) are connected to the origin with lines. Manifold eccentricity of a given region is the length (*i.e.,* Euclidean distance) of this line.
